# Acoustic stimulation increases implicit adaptation in sensorimotor adaptation

**DOI:** 10.1101/2020.10.25.354340

**Authors:** Li-Ann Leow, James R. Tresilian, Aya Uchida, Dirk Koester, Tamara Spingler, Stephan Riek, Welber Marinovic

## Abstract

Sensorimotor adaptation is an important part of our ability to perform novel motor tasks (i.e., learning of motor skills). Efforts to improve adaptation in healthy and clinical patients using non-invasive brain stimulation methods have been hindered by interindividual and intra-individual variability in brain susceptibility to stimulation. Here, we explore unpredictable loud acoustic stimulation as an alternative method of modulating brain excitability to improve sensorimotor adaptation. In two experiments, participants moved a cursor towards targets, and adapted to a 30° rotation of cursor feedback, either with or without unpredictable acoustic stimulation. Acoustic stimulation improved initial adaptation to sensory prediction errors in Study 1, and improved overnight retention of adaptation in Study 2. Unpredictable loud acoustic stimulation might thus be a potent method of modulating sensorimotor adaptation in healthy adults.

## Introduction

A challenge in making accurate movements is the fact that the state of our bodies and our environments often change. For example, we often blink as we move our eyes, perturbing our eye movements. Fatigue changes the way our muscles respond. Sensory feedback of our movements is delayed and corruptible. Despite such noisy, uncertain, and delayed sensory feedback, we adapt rapidly, quickly modifying our movements to attain our goals. This capacity to adapt movements to changes in the body or the environment is termed sensorimotor adaptation, and is an important part of our ability to solve novel motor tasks (i.e., learning of motor skills).

Sensorimotor adaptation is often studied by distorting the relationship between the motor command and the perceived sensory feedback about the movement, for example by disturbing visual feedback of the movement, imposing perturbing forces at the moving limb, or by disturbing acoustic feedback of speech. Such disturbances often evoke discrepancies between predicted and actual sensory consequences of our movements, typically termed sensory prediction errors (Izawa & Shadmehr, 2011). Sensory prediction errors can take the form of discrepancies within sensory modalities (e.g., predicted sensory feedback of a movement does not match perceived sensory feedback of the movement) or between sensory modalities (e.g., visual feedback of hand motion does not match kinaesthetically felt hand movement). Sensory prediction errors can drive a change in the system that predicts sensory consequences of motor commands (Izawa & Shadmehr, 2011). Disturbances also evoke discrepancies between predicted task outcomes and actual task outcomes, here termed task errors. In target-reaching tasks, task errors often take the form of discrepancies between the target position and movement endpoint (Schaefer *et al*., 2012; Gaveau *et al*., 2014; Reichenthal *et al*., 2016), although they can also take the form of failures to achieve more abstract performance goals (Mazzoni & Krakauer, 2006; Taylor & Ivry, 2011). Task errors can elicit the use of explicit strategies (such as volitionally re-aiming to the left of a target to counteract a clockwise rotation of cursor feedback)(Uhlarik, 1973; Mazzoni & Krakauer, 2006), and the formation of associations between cues or stimuli properties within the task context and behavioural responses (Welch, 1971; Cunningham & Welch, 1994; Ishii *et al*., 2018; McDougle & Taylor, 2019; Leow *et al*., 2020).

Standard sensorimotor adaptation paradigms often conflate task errors and sensory prediction errors. However, certain experimental techniques are available to dissociate processes driven by sensory prediction errors and task errors (Welch, 1969; Schaefer *et al*., 2012; Reichenthal *et al*., 2016; Leow *et al*., 2018; Kim *et al*., 2019). These methods offer the opportunity to improve ongoing efforts to improve motor learning in healthy and clinical patients. Such efforts thus far have focussed on non-invasive brain stimulation methods such as transcranial direct current stimulation, with inconsistent results (Jalali *et al*., 2017; Lopez-Alonso *et al*., 2018; Mamlins *et al*., 2019). Importantly, these inconsistent results might result from both insufficient dissociation of mechanisms underpinning adaptation, as well as high inter-individual and intra-individual variability in brain susceptibility to standard non-invasive brain stimulation methods (Chew *et al*., 2015; Labruna *et al*., 2019). It might thus be worth exploring alternative methods of modulating brain excitability to improve motor learning. One promising candidate is loud acoustic stimulation (LAS).

LAS has been used for decades in human motor control studies as a tool to probe movement preparation, providing readouts of the state of the system slightly before a voluntary action is initiated (Carlsen *et al*., 2004; Kumru *et al*., 2006; Forgaard *et al*., 2011; Marinovic *et al*., 2017b). This simple paradigm has very robust behavioural effects such as reduced reaction times and increased response vigour when applied during movement preparation (Valls-Sole *et al*., 1999; Marinovic *et al*., 2013; Castellote & Kofler, 2018; McInnes *et al*., 2020). Such effects are also evident in clinical populations as LAS is capable of facilitating movement initiation and execution in strokes survivors performing constrained or unconstrained multijoint movements (Honeycutt & Perreault, 2012; Honeycutt *et al*., 2015; Marinovic *et al*., 2016; Rahimi & Honeycutt, 2020). Critically, LAS leads to widespread, rapid effects not only on the primary motor cortex (Furubayashi *et al*., 2000; Marinovic *et al*., 2014b), but also in the sensorimotor and executive regions of the brain (Hackley *et al*., 2009; Neuner *et al*., 2010; Marinovic *et al*., 2014a; Chen *et al*., 2016), suggesting it could affect learning processes that depend on these brain regions.

In this study, we explored the possibility that LAS could affect sensorimotor adaptation. In the first study, participants moved a cursor towards targets, and adapted to a 30° rotation of cursor feedback which either induced sensory prediction errors or induced both sensory prediction errors and task errors, using previously validated procedures of dissociating task errors and sensory prediction errors (Leow *et al*., 2018; 2020). LAS was applied randomly in 50% of the adaptation trials. We found that LAS increased adaptation to sensory prediction errors. In the second study, we explored effects of LAS on retention of sensorimotor adaptation, by examining how LAS during initial adaptation to a first perturbation would alter anterograde interference when learning a second, opposite perturbation, after an overnight delay. We found that LAS during initial learning increased anterograde interference of implicit sensorimotor memories, giving support to the idea that LAS increases the persistence of implicit memories formed during sensorimotor adaptation.

## EXPERIMENT 1

### Methods

#### Participants

Sixty-four undergraduate psychology students completed the study in exchange for course credit or monetary reimbursement (mean age =19.9 years, SD =2.7, 45 female). Participants were assigned to the Standard Task Error LAS group (n=16), Standard Task Error no LAS group (n =16), the No Task Error LAS group (n=16) or the No Task Error no LAS group (n =16). Sample size selection was based on similar work employing task error manipulations (Leow *et al*., 2020).

All participants were self-reported to be naïve to visuomotor rotation and force-field adaption tasks. This study was approved by the Human Research Ethics Committee at The University of Queensland. All participants provided written, informed consent.

#### Apparatus and Stimuli

Participants completed the task using a vBOT planar robotic manipulandum, which has a low-mass, two-link carbon fibre arm and measures position with optical encoders sampled at 1,000 Hz (Howard et al., 2009). Participants were seated on a height-adjustable chair at their ideal height for viewing the screen for the duration of the experiment. Visual feedback was presented on a horizontal plane on a 27” LCD computer monitor (ASUS, VG278H, set at 60Hz refresh rate) mounted above the vBOT and projected to the participant via a mirror in a darkened room, preventing direct vision of her/his hand. The mirror allowed the visual feedback of the targets, the start circle, and hand cursor to be presented in the plane of movement, with a black background. The start was aligned approximately 10cm to the right of the participant’s mid-sagittal plane at approximately mid-sternum level. An air-sled was used to support the weight of participants’ right forearms, to reduce possible effects of fatigue.

The LAS stimuli were bursts of 50 ms broadband white-noise with a rise/fall time shorter than 2 ms and peak loudness of 94dBa. Acoustic stimuli were presented through high fidelity stereophonic headphones (Seinheiser model HD25-1 II; frequency response 16Hz to 22kHz; Sennheiser Electronics GmbH & Co. KG, Wedemark, Germany). The LAS intensity choice was based on the knowledge that intensities above 79dBa can reliably change cortical excitability within the primary motor cortex (Furubayashi *et al*., 2000).

#### Trial Structure

While grasping the robot arm, participants moved their hand-cursor (0.5cm radius red circle) from the central start circle (0.5cm radius white circle) to the targets (0.5cm radius yellow circles). Targets appeared in random order at one of eight locations (0°, 45°…. 315°) at a radius of 9 cm from a central start circle. At the start of each trial, the central start circle was displayed. If participants failed to move their hand-cursor to within 1cm of the start circle after 1 second, the robotic manipulandum moved the participant’s hand to the start circle (using a simulated 2-dimensional spring with the spring constant magnitude increasing linearly over time). A trial was initiated when the cursor remained within the home location at a speed below 0.1cm/s for 200ms.

Across all experiments, we used a classical timed-response paradigm (Schouten & Bekker, 1967) to manipulate movement preparation time during the planar reaching task (Favilla & De Cecco, 1996; Marinovic *et al*., 2017a). A sequence of three tones, spaced 500ms apart, was presented at a clearly audible volume via external speakers. Participants were instructed to time the onset of their movements with the onset of the third tone, which was more highly-pitched than the two previous, and then to slice through the target with their cursor. Movement initiation time was identified online as when hand speed exceeded 2cm/s.

To familiarize participants with the equipment and the timed-response paradigm, all participants were first allowed a familiarization block comprising a maximum of 6 cycles. One cycle consisted of 1 trial to each of the 8 targets, and target order was random within each cycle. Participants were explicitly instructed to make movements to slice through the targets, rather than to stop on the targets.

After familiarisation, all participants (regardless of assigned condition) were given the same task instruction, as follows. “Your task in this experiment is to hit the targets. The computer might disturb the cursor and/or the target, this is a normal part of the experiment, just try to hit the target as well as you can”. Participants then completed the following blocks in sequence. **<u>Baseline block</u>** (6 cycles): no rotation of visual feedback. **<u>Training block</u>** (60 cycles): During training, half of all participants encountered a clockwise 30°cursor rotation and half encountered a 30° counterclockwise cursor rotation, such that rotation direction was counterbalanced for all conditions. **Task error manipulations:** The standard task error groups experienced standard task errors, where the target remained stationary throughout the trial, such that the perturbation evoked a task error (failure to attain the target) (see Figure 1 bottom). The no task error groups experienced the no task error manipulation, by moving the target mid-movement to align to the cursor direction when the cursor reached 4cm of the 9cm distance between the start and the target (see Figure 1 top). This is analogous to moving a basketball hoop towards the ball mid-flight—the ball always goes through the hoop regardless of the participant’s actions). **LAS manipulations:** Upon movement completion, the LAS groups (i.e., half of the standard task error group (n=16) and half of the no task error group (n=16)) received LAS upon movement completion in a randomly selected 50% of trials (i.e., 4 out of 8 trials in every cycle). No LAS was imposed for the noLAS groups (half of the standard task error group (n=16) and half of the no task error group (n=16)). Task error and LAS manipulations were restricted to the training block. **Instructions of perturbation removal:** Upon completing the adaptation block, all participants received explicit instructions about the rotation removal, as follows: “Any disturbance that the computer has applied is now gone, and the feedback of your movement will now be hidden as soon as it leaves the start circle, so please move straight to the target”. **<u>No feedback block</u>** (6 cycles): Upon leaving the start circle, no feedback about movements was available. Reaches that remain adapted despite explicit knowledge of perturbation removal indicate implicit aftereffects. **<u>Washout block</u>** (12 cycles): Cursor position feedback was restored, but the 30° rotation of cursor (or target) was removed. In the baseline, adaptation, and the washout trials, participants received cursor feedback as soon as the cursor travelled outside the 0.5cm start circle, and terminated after the cursor travelled outside the 9cm radius between the start and the target.

**Figure 1.**
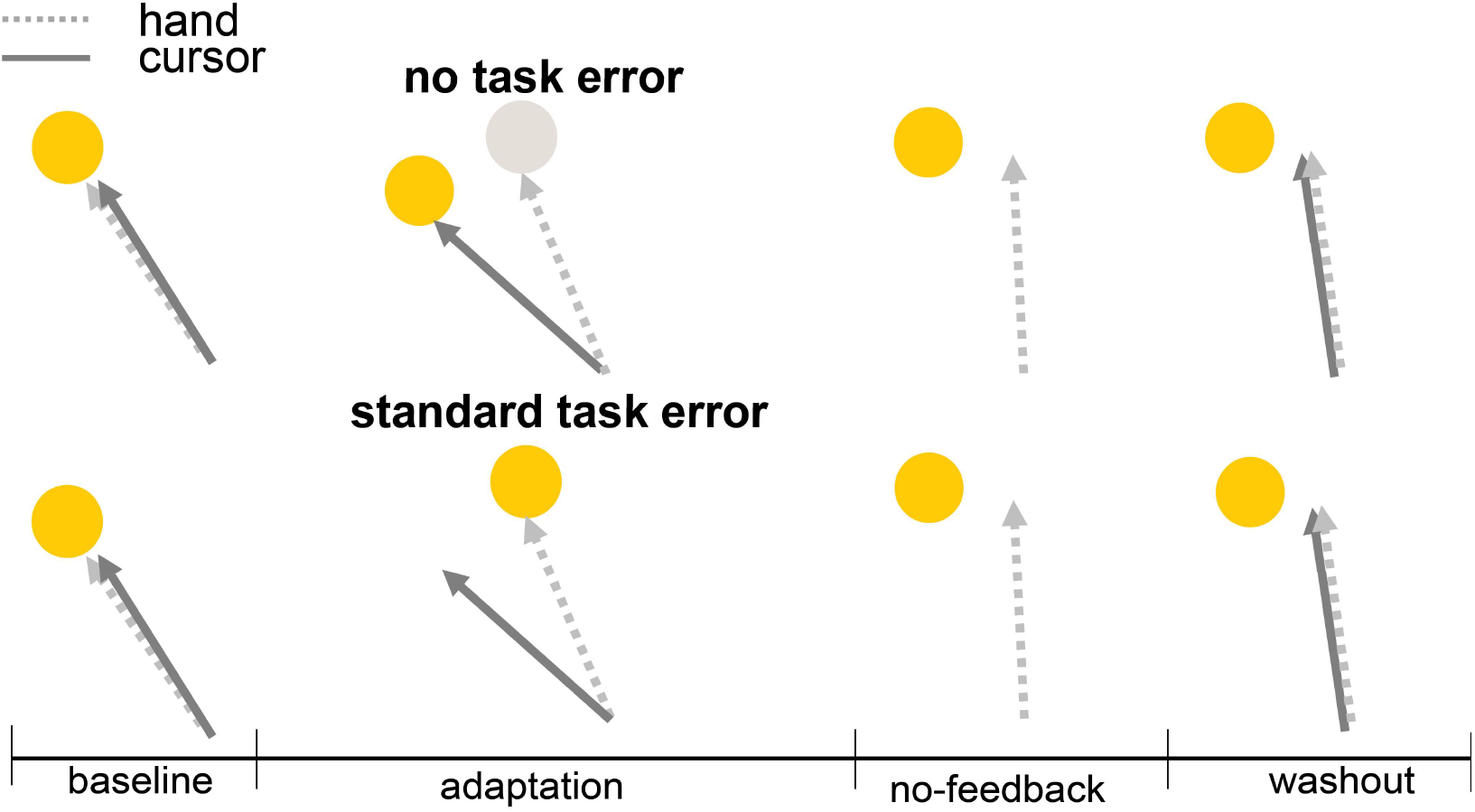
Study 1 protocol, where participants completed baseline (6 cycles with veridical cursor feedback of hand position), adaptation (60 cycles with rotated cursor feedback, with either no-task errors or standard task errors, and LAS applied in every 4 out of 8-trial cycle), no-feedback (6 cycles with no cursor feedback), and washout (12 cycles with veridical cursor feedeabk).

#### Data analyses

For all blocks, the estimated adaptation performance was estimated as percent adaptation, which estimates reach directions relative to the ideal reach direction.

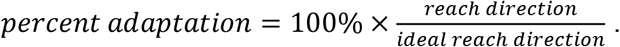

Movement direction was measured at 25% of the movement distance after the cursor left the home position, similar to our previous work (Leow *et al*., 2018).

Unlike p-values, Bayes factors do not have the tendency to over-estimate the evidence against the null hypothesis (Gelman & Tuerlinckx, 2000; Wetzels et al., 2011). We thus chose to use Bayesian statistics to evaluate evidence for the alternative hypothesis and for the null hypothesis. Analyses were conducted in JASP (Team, 2020). The default Cauchy prior widths (0.707) values in JASP were used to quantify the relative evidence that the data came from the alternative versus a null model. Jeffreys’s evidence categories for interpretation (Wetzels et al., 2011), were taken as the standard for evaluation of the reported Bayes Factors.

##### Experiment 1

Bayesian ANCOVAs were run with between-subjects factors LAS (LAS, noLAS) and Task Error (No Task Error, Task Error), and pre-rotation biases entered as covariates of no interest. Where applicable, within-subjects factors of Phase (Early, Middle, Late) x Cycles (1.20) were entered. Specifically, Bayes factors were used to evaluate main effects, by comparing a model that includes the factor LAS (noLAS, LAS) to a null model without the factor LAS, and a model that includes the factor Task Error (NoTaskError, TaskError) to a null model without the factor Task Error. To quantify the relative evidence for the presence of an interaction effect, compared to its absence, we compared a model which incorporates the interaction between LAS and TaskError to a model that includes only the main effects. For these analyses, the Bayesian inclusion factor is reported (BFinclusion). Post-hoc comparisons corrects for multiple testing by fixing to 0.5 the prior probability that the null hypothesis holds across all comparisons (Westfall *et al*., 1997). Individual comparisons are based on the default t-test with a Cauchy (0, r = 1/sqrt(2)) prior.

### Results

#### Adaptation

Replicating previous results (Leow *et al*., 2018), removing task errors resulted in reduced adaptation than standard task errors (see Figure 2), BF incl = ∞. There was an LAS x Task Error interaction (BF incl = 42375.023), and an LAS x Task Error x Phase interaction, (BF incl = 180062.854). Follow-up LAS x Phase x Cycles Bayesian ANCOVAs were run separately for the no task error and standard task error conditions. There was strong evidence for the main effect of LAS for the no task error group (BF incl = 86209.711, post-hoc Bayesian t-tests: BF10 =6229.14), but weak evidence for the main effect of LAS for the standard task error group (BF incl = 0.193).

**Figure 2.**
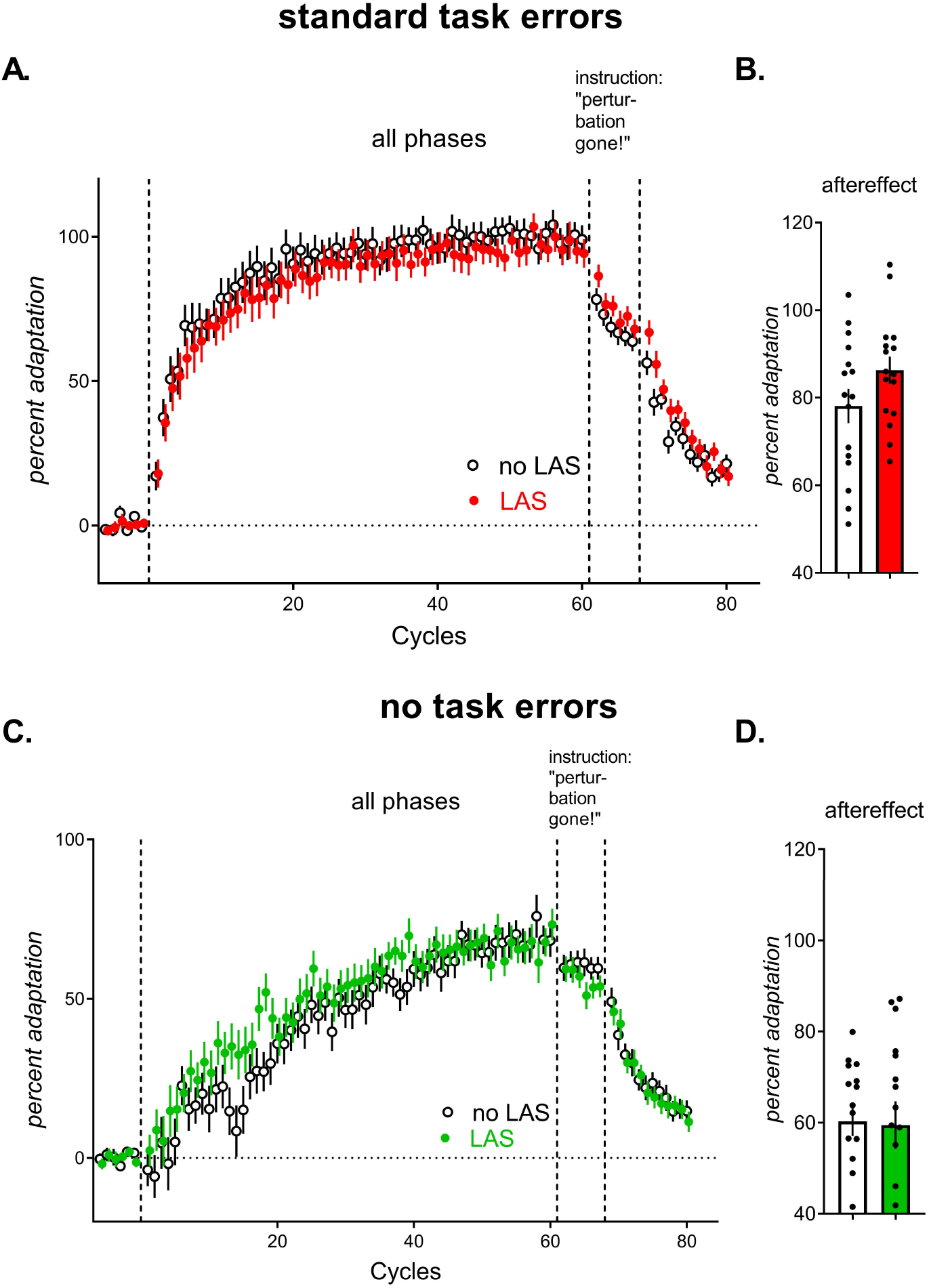
Cycle-by-cycle percent adaptation, in the standard task error groups (A), and the no-task error groups (C). B&D show percent adaptation in the implicit aftereffect measured in the first no-feedback cycle. Error bars are SEM.

#### Implicit aftereffects

Reaches that remain adapted despite explicit knowledge that the perturbation is absent are suggestive of a remapping of the relationship between motor commands and the predicted sensory feedback of the ensuing movement, termed implicit aftereffects. Implicit aftereffects were larger with standard task errors than with no task errors (Figure 1B&F), replicating previous results (Schaefer *et al*., 2012; Leow *et al*., 2018; Kim *et al*., 2019) (main effect of task error BF incl = 21519.029). The evidence for an LAS effect on implicit aftereffect was weak (LAS main effect, BF incl = 0.376), regardless of task error (LAS x Task Error interaction, BF incl = 0.556).

#### Washout

Returning visual feedback to the veridical state in the washout phase to measure aftereffects reflects persistence of sensorimotor adaptation, as well as active compensation for the error that results from abrupt removal of the cursor rotation. As expected, movements in the washout phase quickly decayed to the unadapted state across all conditions (of the washout block with LAS, which decayed at the later cycles to be similar to the noLAS group (Figure 2). The evidence for an LAS effect on washout was weak (LAS main effect, BF incl = 0.357), regardless of task error (LAS x Task Error interaction, BF incl = 0.448).

### Experiment 1 summary

In the absence of task errors, LAS increased percent adaptation, indicating that LAS boosted adaptation to sensory prediction errors. In the presence of task errors, LAS showed no effect of LAS. Thus, in the presence of task errors, the effect of LAS on sensory prediction errors might have been masked by components of adaptation driven by task errors.

## EXPERIMENT 2

We extended Experiment 1 by additionally testing whether if LAS during adaptation to a first perturbation would be evident when adapting to a second, directionally opposite perturbation after an overnight period. Without the passage of time between learning the first and the second perturbation, this protocol typically results in interference in adaptation to the first perturbation to adaptation to the second perturbation (Brashers-Krug *et al*., 1996; Sing & Smith, 2010; Huang *et al*., 2011; Leow *et al*., 2013; Herzfeld *et al*., 2014; Leow *et al*., 2014; Leow *et al*., 2016; Maeda *et al*., 2018; Lerner *et al*., 2020). This anterograde interference can dissipate after washout and with the passage of time (Krakauer *et al*., 2005), although this pattern of time-dependent reduction in anterograde interference is not evident in all studies (Goedert & Willingham, 2002; Caithness *et al*., 2004; Cothros *et al*., 2006).

### Methods

#### Participants

Fifty undergraduate psychology students participated in the study for course credit (mean age = 20.1, range = 17 - 30 years, 27 women, 2 left-handed). Four participants did not return for part 2. These four incomplete datasets were discarded. Participants were assigned to the LAS condition (n=16), the noLAS condition (n=15), or the naïve control condition (n=15). All participants were naïve to visuomotor rotation and force-field adaption tasks. This study was approved by the Human Research Ethics Committee at The University of Queensland. All participants provided written, informed consent.

#### Apparatus and stimuli

Due to limited availability of the vBOT at the time, we conducted Experiment 2 on a WACOM digitizing tablet, which is a validated mode of data collection for visuomotor adaptation experiments. Participants controlled an on-screen cursor by moving a digitizing stylus with their right-hand on a digitizing tablet (WACOM Intuos4 PTK 1240, size: 19.2 x 12 in., resolution ± 0.25 mm), that recorded X-Y coordinates of the stylus position approximately every 10ms. The tablet and the monitor were both mounted horizontally such that a planar movement on the tablet translated to a planar on-screen movement. The computer monitor (60Hz) was mounted 28 cm mounted above the tablet. Participants were seated on a chair at their ideal height for viewing the computer screen for the duration of the experiment.

Throughout the experiment, participants wore high fidelity stereophonic headphones, which presented all acoustic stimuli (Steinheiser model HD25-1 II; frequency response 16 Hz to 22 kHz; Sennheiser Electronics GmbH & Co. KG, Wedemark, Germany). Sound intensity was measured with a DIGITECH sound level meter with calibrator (model: QM 1592, A & C weighted; Digitech, Sanda, Utah, United States) placed 2 cm from the headphone speaker.

LAS stimuli were bursts of 50 ms broadband white-noise with a rise/fall time shorter than 2 ms and peak loudness of 80 dBa.

#### Trial structure

During the task, participants made centre-out reaching movement by moving the pen from a green central home position (8 pixels) to a red target (15 pixels). Feedback of the position of the stylus was indicated by a small black cursor (5 pixels). Participants had to move the cursor to the home and stay there for 1 second before the target appeared. A soft beep (maximum 60dB) sounded in conjunction with target appearance, signalling participants to move to the target. Targets appeared at one of eight locations that were separated by 45° with a radius of 7 cm (0°, 45°…315°), presented in random order. Participants were instructed to “not to stop on the target, but to slice through it and stop afterwards, before you go back to the centre, and make sure you move fast.” Movement completion was defined as the time when the displacement from the centre of the start circle was greater than 280 pixels. (i.e., as soon as the XY coordinates stopped changing) and the pen had moved a minimum of 280 pixels on the tablet. Upon movement completion, a second beep sounded, signalling participants to return the cursor to the start. To assist with this, the cursor distance from the target was represented by a black ring with a radius equal to the distance of the pen from the start position. As the pen position got closer to the start, the circle shrank. The actual cursor position was shown when 60 pixels away from the centre of the start.

All Experiment 2 participants completed the following blocks in sequence (see Figure 3), where one cycle consisted of one trial to each of the eight targets: **<u>Baseline block</u>** (15 cycles): no rotation of visual feedback. **<u>Adaptation block A</u>** (30 cycles): For the LAS group (n=16) and the noLAS group (n=16), cursor feedback was rotated by 30° clockwise. The target did not move during the trial, and thus the rotation resulted in standard task errors (missing the target). For the naïve group (n=15), no cursor rotation was imposed. In randomly selected 50% of trials (i.e., 4 out of 8 trials in each cycle, LAS was imposed immediately after movement completion for the LAS group, but not for the noLAS or the naïve group). LAS was never employed in any other block other than A1. **Instruction of perturbation removal**: an on-screen popup with the following statement appeared: “Now, all disturbances that the computer applied will be removed. We will also hide feedback of the cursor as soon as it leaves the start. Just move straight to the target.” This popup was read aloud by the experimenter to ensure participant comprehension. **<u>No-feedback block A</u>** (1 cycle): In the no-feedback cycle block, cursor feedback was removed as soon as it left the start circle: reaches that remain adapted despite explicit knowledge of perturbation removal indicate implicit aftereffects. **<u>Washout block with no task errors</u>** (30 cycles): cursor feedback was returned, but cursor rotation was absent. Error upon perturbation removal are known to improve subsequent adaptation, particularly if the errors upon perturbation removal are of the same sign as that experienced in subsequent perturbation (Herzfeld, Vaswani, Marko, & Shadmehr, 2015). It is unclear however if task errors contribute to this effect (Orban de Xivry & Lefèvre, 2015). We thus employed the no-task-error manipulation, removing task error by moving the target 150ms into the movement to align with the on-screen cursor direction (Leow et al., 2018). **Delay:** Participants left the lab for a delay between 17h to 7 days 4h (mean: 2d 7h). **<u>Adaptation block B</u>** (30 cycles): The cursor was rotated 30° in the opposite direction to that experienced on day 1 (counterclockwise), with the no-taskerror manipulation described above. **<u>No-feedback block B</u>** (30 cycles): Before starting the no-feedback block, participants were explicitly instructed that the rotation was removed by an on-screen popup with the same instructions as for the No-feedback block A. We chose 30 cycles instead of 1 cycle to additionally test for LAS effects on the persistence of implicit aftereffects across increasing no-feedback cycles.

**Figure 3.**
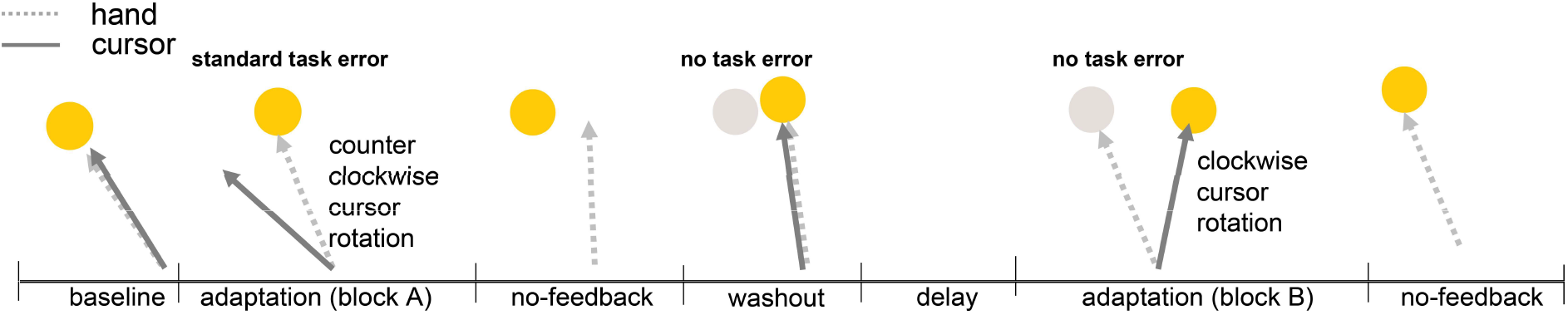
Study 2 protocol, where participants completed baseline (15 cycles with veridical cursor feedback of hand position), adaptation block A (30 cycles with counterclockwise cursor rotation with standard task errors, and LAS applied in a randomly selected 4 out of 8-trials in a cycle), no-feedback (1 cycle with no cursor feedback), and washout (30 cycles with veridical cursor feedback, no task error). After an overnight delay, participants encountered block B where they adapted to a clockwise cursor rotation with no task errors, followed by a 30-cycle no-feedback block. The naïve control group was the same as the LAS and no-LAS groups, except that they had no cursor rotation (veridical cursor feedback) in adaptation block A.

#### Data analysis

To test how the LAS manipulation altered adaptation, Bayesian ANCOVAs were run with between-subjects factors Condition (naïve, LAS, noLAS) and pre-rotation biases as covariates (biases estimated from the mean percent adaptation from the last 3 cycles before rotation onset). Where applicable, the within-subjects factors phase (early, late) and cycles (cycle 1.15) were entered. Analyses on Day 1 did not include data from the naïve control group, who, unlike the LAS and no-LAS group, did not experience any rotation on Day 1.

### Results

After an overnight delay at least 17 hours, participants adapted to an opposite perturbation (cursor feedback was rotated 30°) than on day 1. Reach directions averaged across every cycle is shown in Figure 4 (day 1) and Figure 5 (day 2), where data from LAS group are shown in red, data from the noLAS group are shown in clear circles, and data from the naïve group are shown in blue boxes.

**Figure 4.**
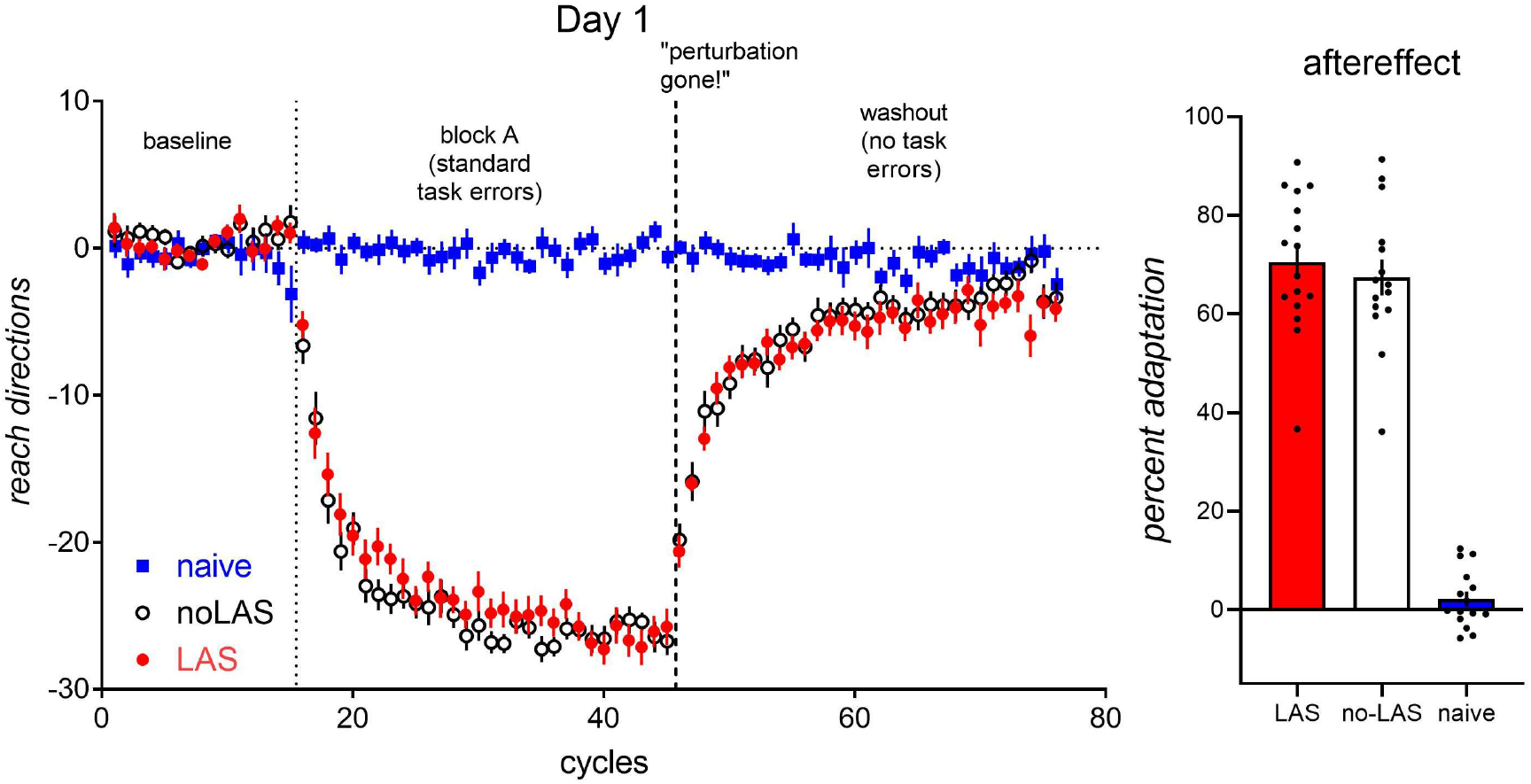
Left: Day 1 reach directions. All groups experienced 15 cycles at baseline without rotation followed by 30 adaptation cycles (30° clockwise rotation for the noLAS and LAS group, no rotation for the naïve control group), one no-feedback cycle and 30 washout cycles with no task errors. Right: aftereffects measured after explicit notification that the perturbation had been removed. All error bars are SEM.

**Figure 5:**
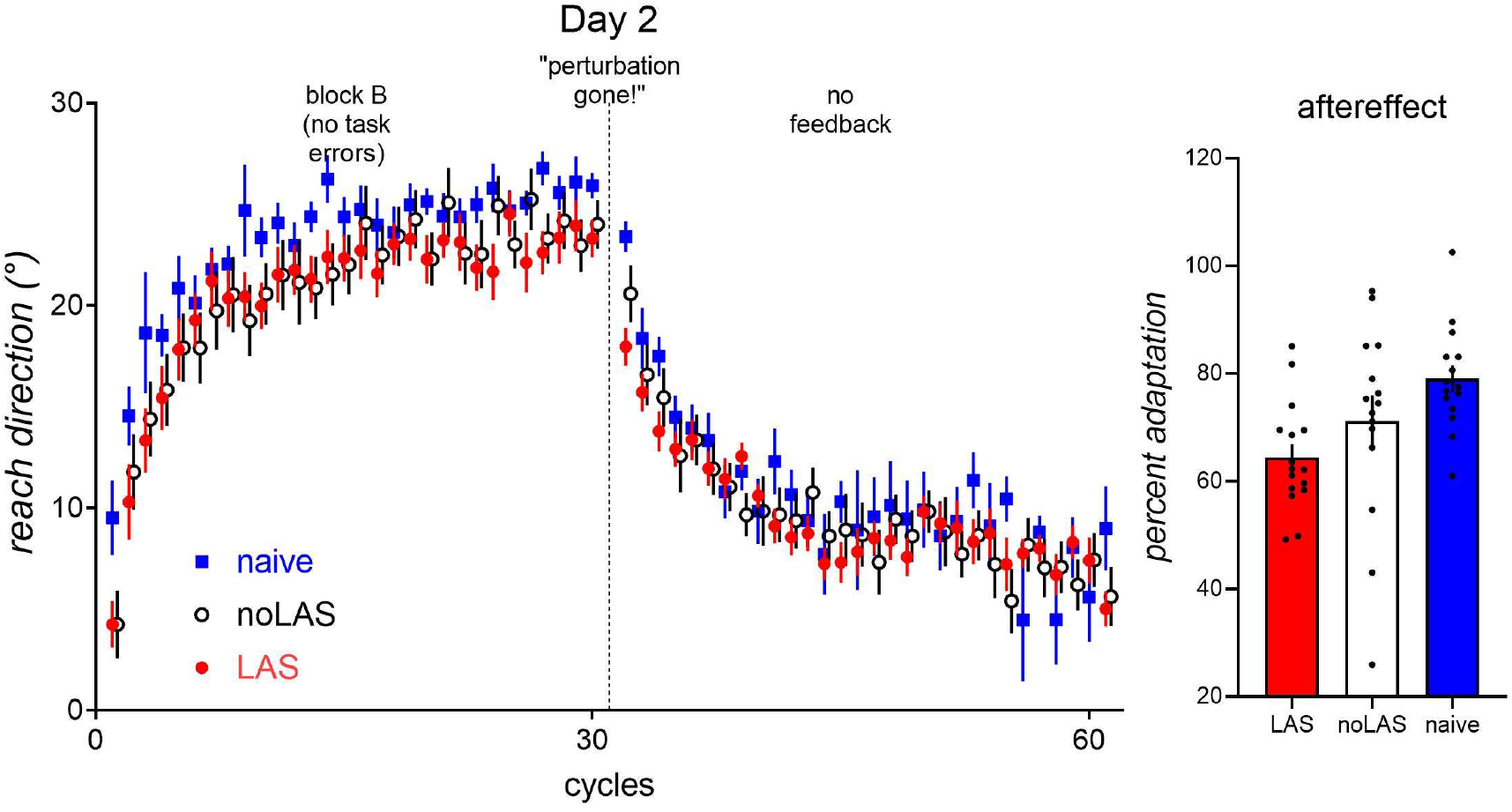
Left Day 2 reach directions. All groups were exposed to a 30° counterclockwise rotation in B with no task errors, followed by 30 no-feedback cycles. Right: implicit aftereffect measured after explicit notification that the perturbation had been removed. Error bars are SEM.

#### Day 1 Adaptation

In block A, participants adapted to a 30° counterclockwise perturbation with standard task errors (discrepancies between desired task outcomes and actual task outcomes) either with LAS (LAS) or no LAS (noLAS) on day 1, or did not adapt to a perturbation (naïve control). Day 1 adaptation was thus similar to the standard task error group in Experiment 1. Figure 4 (left) shows cycle-by-cycle adaptation performance in the LAS and noLAS groups. To examine whether LAS altered adaptation in the presence of standard task errors on Day 1, we ran LAS (LAS, noLAS) x Phase (early, late) x Cycles (Cycle 1.15) Bayesian ANCOVAs on the adaptation block where the covariate was the prerotation percent adaptation. Replicating results from the standard task error group in Experiment 1, LAS did not affect adaptation (main effect of LAS: BF incl = 0.261, all interactions with LAS BF, incl < 1).

#### Day 1 implicit aftereffects

Replicating Experiment 1, the size of the implicit aftereffect measured in the first no-feedback cycle, did not differ between the LAS and the noLAS group (see Figure 4 right), as one-way Bayesian ANCOVA with the factor LAS (no-LAS, LAS) with pre-rotation bias as covariate showed weak-to-little evidence for any LAS effect, BF incl = 0.393.

#### Day 1 washout

The main effect of LAS, and all interactions with LAS showed little-to-weak evidence of any effect of LAS on the post-adaptation washout phase (BF incl < 1).

#### Day 2 Adaptation

After Day 1 washout, all participants had an overnight delay (>17 hours), and returned on day 2 to adapt to a directionally opposite perturbation (i.e., B) with no task error (see Figure 5). A one-way Bayesian ANCOVA with the factor Condition (LAS, noLAS, naïve) and covariate (pre-rotation bias) was run to assess the state of adaptation upon returning after the overnight delay. Groups differed on the first cycle of block B, (main effect of Group BF incl = 4.372). Post-hoc tests showed that the LAS group was worse-than-naïve, BF10 = 5.826. In contrast, the No-LAS group was not worse-than-naïve, BF10 = 2.064. Thus, the overnight delay and washout removed anterograde interference effects for the first block B cycle only for the no-LAS group: the LAS group showed persistent anterograde interference effects in the first B cycle.

To examine whether LAS altered adaptation on day 2, we ran Condition (LAS, noLAS, naïve) x Phase (early, late) x Cycles (Cycle 1.15) Bayesian mixed ANCOVAs on adaptation block B on day 2. Groups did not differ with/without LAS or in comparison to naïve (see Figure 5 left) [main effect of Condition, BF incl = 0.966, Phase x Condition, BF incl = 2.144, Phase x Cycles x Condition interaction, BF incl = 2.373].

#### Implicit aftereffects

Worse-than-naive implicit adaptation to B on Day 2 is indicative of persistent anterograde interference effects from implicit adaptation to the Day 1 perturbation (see Figure 4 right). One-way Bayesian ANCOVA showed a main effect of Condition, BF incl = 10.697, as the LAS group showed smaller implicit aftereffects than naïve BF 10 = 455.097. In contrast, the no-LAS group tended to not differ from naïve (BF 10 = 0.916). The LAS and no-LAS groups however, did not appear to differ from each other (BF10 = 1.135).

### Experiment 2 summary

LAS on Day 1 resulted in worse-than-naive initial adaptation to an opposite rotation on Day 2, and worse-than-naïve implicit aftereffects on Day 2. Thus, anterograde interference from Day 1 to Day 2 was present in the LAS group. In contrast, such anterograde interference was reduced with washout and the passage of time for the noLAS group, as adaptation and implicit aftereffects did not differ from naive. Thus, for the LAS group, the washout and subsequent overnight delay failed to remove anterograde interference. Implicit aftereffects do not arise in the absence of sensory prediction errors (Izawa & Shadmehr, 2011), and thus aftereffects that persist despite explicit notification of perturbation removal are thought to be a reasonable measure of implicit adaptation to sensory prediction errors (Maresch & Donchin, 2019). We interpret the anterograde interference in implicit aftereffects in the LAS group to suggest that LAS on Day 1 resulted in a more robust adaptation to sensory prediction errors on Day 2.

## Discussion

There is a growing literature on the effects of acoustic stimulation on movement initiation and execution — reduced onset latency and increase response vigour — when the LAS is presented during movement preparation (for reviews see (Valls-Sole, 2012; Nonnekes *et al*., 2015; Marinovic & Tresilian, 2016). These robust effects of LAS, however, have been observed only immediately, affecting the upcoming response, rather than on subsequent movements. The experiments reported here sought to determine whether random presentation of LAS during adaptation of reaching movements to rotated movement feedback, at movement completion, could regulate the extent of implicit adaptation when LAS presentation ceased.

Experiment 1 explored if LAS would affect sensorimotor adaptation with sensory prediction errors alone, or with both task errors and sensory prediction errors. We found that LAS boosted adaptation to sensory prediction errors under no-task-error conditions, particularly at initial exposure to the perturbation. Under standard task error conditions, this effect of LAS might have been masked by task-error driven components of adaptation (such as explicit re-aiming strategies (Mazzoni & Krakauer, 2006) or stimulus-response associations) (Ishii *et al*., 2018; McDougle & Taylor, 2019; Leow *et al*., 2020), as it was not discernible under standard task error conditions. In Experiment 2, we further explored the capacity of LAS to influence the retention of sensorimotor memories. Retention was examined via anterograde interference: where persistent memories acquired from adaptation to an initial perturbation interferes with subsequent adaptation to a second, different perturbation (Brashers-Krug *et al*., 1996; Shadmehr & Brashers-Krug, 1997; Caithness *et al*., 2004; Miall *et al*., 2004; Krakauer *et al*., 2005; Cothros *et al*., 2006; Sing & Smith, 2010; Leow *et al*., 2013; Lerner *et al*., 2019). In Experiment 2, we presented either LAS or no LAS during adaptation to a counterclockwise rotation of cursor feedback under standard task error conditions on the first day. Replicating Experiment 1, we did not find an effect of LAS under standard task error conditions on the first day. On the second day, all participants were exposed to a clockwise rotation under no-task-error conditions, in the absence of any LAS. Participants exposed to LAS on the first day showed worse-than-naïve adaptation, in contrast to the no-LAS group who did not differ from naïve. Furthermore, the LAS group also showed worse-than-naïve Day 2 aftereffects, suggesting robust anterograde interference in aftereffects in the LAS group. In contrast, aftereffects in the no-LAS group did not differ reliably from naïve, suggesting a time-dependent decay of anterograde interference effects in the no-LAS group. Thus, LAS at initial learning seemed to have boosted the retention of sensorimotor memory, such that it increased subsequent anterograde interference on implicit aftereffects on Day 2. Taken together, Experiment 1 and 2 results suggest that LAS during exposure to a sensorimotor perturbation boosts the acquisition and retention of implicit adaptation to perturbation-induced sensory prediction errors.

The results of our experiments are broadly in agreement with the literature on the effects of physiological arousal on memory formation (Schwarze *et al*., 2012; McGaugh, 2018). Intense, unpredictable events such as LAS are powerful modulators of arousal, which can enhance memory for goal-relevant stimuli in many cognitive domains (Mather & Sutherland, 2011; Mather *et al*., 2016; Clewett *et al*., 2018). Salient stimuli activate the locus coeruleus, resulting in quick, widespread release of noradrenaline in many regions of central nervous system (Brun *et al*., 1993; Joshi *et al*., 2016). Noradrenaline has been shown to increase the signal-to-noise ratio, improving synaptic transmission, of both excitatory and inhibitory afferents (Woodward *et al*., 1991; Jiang *et al*., 1996; Morilak *et al*., 2005). This improvement in synaptic transmission is broadly consistent with the pattern of improved implicit memory formation with LAS in our study. In what follows, we discuss possible physiological mechanisms by which LAS could interact with error processing mechanisms in our paradigm.

Sensorimotor adaptation involves plastic changes in the cerebellum (Martin *et al*., 1996; Tseng *et al*., 2007; Therrien *et al*., 2016), posterior parietal cortex (Diedrichsen *et al*., 2005), the somatosensory cortex (Bernardi *et al*., 2015; Ohashi *et al*., 2019), the primary motor cortex (Cothros *et al*., 2006; Richardson *et al*., 2006; Hadipour-Niktarash *et al*., 2007; Galea *et al*., 2011; Kawai *et al*., 2015; Perich *et al*., 2017), and the basal ganglia (Shadmehr & Holcomb, 1999), for reviews, see (Shadmehr & Krakauer, 2008; Haar & Donchin, 2019). In particular, neurons of the primary motor cortex appear to encode errors immediately post-movement: stimulation of these neurons at 100ms postmovement but not after 100ms produced adapted behaviour: this stimulation induced adaptation increased gradually and decreased gradually upon cessation of stimulation, similar to actual sensorimotor adaptation (Inoue *et al*., 2016). One hypothesis is that premotor/M1 neurons are engaged by and encode errors with respect to their preferred direction, which in turn engages the cerebellar Purkinje cells, and drive a change in the outgoing motor command (Inoue *et al*., 2016). Repeated exposure to such errors might alter directional selectivity of parietal cortex neurons (Haar *et al*., 2015; Inoue & Kitazawa, 2018), as well as changes in functional connectivity between the parietal and motor cortices (Tanaka *et al*., 2009). At this stage we do not know whether LAS exerts its effects on adaptation by affecting function in M1, the cerebellum, parietal cortex, or the somatosensory cortex. LAS may well affect activity in many or all of these regions; for example, salient events are known to increase complex spiking activity in cerebellar Purkinje cells (Heffley *et al*., 2018). We speculate that salient, intense events such as LAS presented in conjunction with sensory prediction errors increases the sensitivity of M1 neurons to sensory prediction errors. It is known that LAS can rapidly (~ 50 ms) and transiently affect excitability within M1 (Furubayashi *et al*., 2000; Ilic *et al*., 2011; Marinovic *et al*., 2014b; Chen *et al*., 2016). Importantly, the effect of LAS on M1 occurs within the same 100ms timeframe post-movement as when error processing is thought to occur in M1 (Inoue *et al*., 2016). The nature of these rapid changes in M1 excitability depends on the state of movement for action: LAS effects on M1 are excitatory during movement preparation (Marinovic *et al*., 2014b), and inhibitory at other times (Furubayashi *et al*., 2000; Marinovic *et al*., 2014b; Chen *et al*., 2016). If LAS-induced excitatory effects on M1 is the mechanism by which stronger implicit learning is achieved, our results suggest that the effect of LAS on M1 at movement completion should be excitatory, improving M1 excitability during memory formation in visuomotor adaptation paradigms (Hadipour-Niktarash *et al*., 2007; Hamel *et al*., 2017; Spampinato *et al*., 2019). It remains to be tested whether LAS at movement completion excites or inhibits M1 neurons. However, the fact that movement preparation levels ought to be low at movement completion would suggest that the rapid effects of post-movement LAS on M1 would be inhibitory rather than excitatory.

As the locus coeruleus is the main site of norepinephrine production in the central nervous system and responds promptly to salient sensory events, future studies may examine whether differences in sensorimotor memory formation and retention can be predicted by its phasic activation. This could be achieved by recording pupillary changes as a surrogate measure for locus coeruleus activation (Joshi *et al*., 2016). Future studies should investigate whether there is an association between pupillary changes induced by movement execution (Kalwani *et al*., 2014) or salient events (Experiment 1 and 2) and the extent of implicit adaptation to sensory prediction errors across individuals in motor learning. Previous research using a predictive-inference task showed that task-independent changes in pupil dilation, induced by switching the auditory cue at trial onset, led to systematic performance changes across individuals that were dependent on baseline pupil diameter (Nassar *et al*., 2012). In agreement with our findings, these results suggest that well-timed, brief auditory stimuli can alter the state of the central nervous system and affect human performance. We note however that although we suggested a role for noradrenaline in LAS effects of adaptation, we cannot rule out a role for other neurotransmitters. Indeed, there is consensus that dopamine plays a role in processing salient events (Horvitz, 2000; Bromberg-Martin *et al*., 2010): for example, dopamine manipulations modulate midbrain responses to unexpected sounds but not to expected sounds (Valdes-Baizabal *et al*., 2020). Similar to noradrenaline, dopamine acts as a neuromodulator that cannot directly excite or inhibit postsynaptic responses, but modulate the postsynaptic responses to other neurotransmitters. Both noradrenaline and dopamine might thus act to enhance neural gain (Servan-Schreiber *et al*., 1990), increasing sensitivity to sensory prediction errors. Future studies can employ pharmacological manipulations of dopamine and noradrenaline to clarify how noradrenaline or dopamine-dependent processes affect the way LAS alters sensorimotor adaptation.

Although we proposed that the effects of LAS resulted from arousal-induced modulation of sensorimotor adaptation, an alternative, related interpretation is that LAS served as a form of unconditioned stimulus. Loud abrupt bursts of white noise are highly effective unconditioned stimulus in classical conditioning experiments (Morris *et al*., 1998). Indeed, white noise bursts as unconditioned stimuli are more reliable, more extinction-resistant, and result in more stable conditioning compared to electric shocks in fear conditioning experiments (Sperl *et al*., 2016). The finding of day 1 LAS improving day 2 retention in Experiment 2 is broadly consistent with a partial-reinforcement extinction effect, where partial reinforcement schedules improve retention compared to continuous reinforcement schedules (Jenkins & Stanley, 1950). This interpretation does not, however, fully explain the faster adaptation with LAS shown in Experiment 1 under the no-task-error conditions, as it is unclear why participants would adapt more quickly, as faster adaptation does not change the occurrence of the loud acoustic stimulation. If LAS did, indeed, act as a form of unconditioned stimulus, then LAS would have weaker effects if presented on a continuous reinforcement schedule than on a partial reinforcement schedule: this idea remains to be tested experimentally.

Another interpretation is that LAS acted as a form of punishment. Sensorimotor adaptation can be sped up by punishment via monetary losses, both when losses were not task contingent (Song & Smiley-Oyen, 2017), as well as when losses were task contingent (Galea *et al*., 2015; Huang *et al*., 2018b; Song *et al*., 2020), although effects of punishment on adaptation have been inconsistent, with studies showing either similar effects of rewards and punishment (Quattrocchi *et al*., 2017), or no effect of punishment on adaptation rate (Huang *et al*., 2018a; Quattrocchi *et al*., 2018a; Hill *et al*., 2020). Relevant to the current findings however is the fact that in the majority of published studies show no effect of punishment on post-adaptation retention (Galea *et al*., 2015; Song & Smiley-Oyen, 2017; Huang *et al*., 2018b; Quattrocchi *et al*., 2018b), in contrast to the effect of LAS improving post-adaptation retention on Day 2 shown here. Thus, our findings appear inconsistent with the interpretation that LAS acted as a form of punishment during sensorimotor adaptation.

## Conclusion

To summarize, Experiment 1 shows that pairing uncertain, intense auditory stimuli with exposure to a cursor rotation increases adaptation to sensory prediction errors. Experiment 2 further shows that pairing uncertain, intense auditory stimuli has long-lasting effects evident on the next day, manifested in worse-than-naïve adaptation upon initial exposure to a different perturbation, and worse-than-naïve aftereffects that persisted despite explicit knowledge of perturbation removal. Our results provide the first evidence that task-irrelevant loud acoustic stimulation can improve implicit memory formation during adaptation to movement perturbations. We speculate that our findings could be explained by the release of norepinephrine and/or dopamine throughout the central nervous system following the presentation of the LAS, particularly affecting error processing within the primary motor cortex. The results also suggest that unpredictable acoustic stimulation might be a promising way of modulating motor learning in healthy and clinical patients.

Li-Ann Leow: conceptualization, data analysis, writing - original draft, review & editing. James R. Tresilian: writing - review & editing, funding acquisition. Aya Uchida: data collection, project administration, writing - review & editing. Dirk Koester: writing - review & editing. Tamara Spingler: data collection, project administration, writing - review & editing. Stephan Riek: funding acquisition, writing - review. Welber Marinovic: funding acquisition, conceptualization, writing - original draft, review & editing.

## Acknowledgements

This work was funded by an Australian Research Council Discovery Project Grant (DP16012001) awarded to Welber Marinovic, James Tresilian, and Stephan Riek, a UQ Development Fellowship UQFEL1718737 awarded to Li-Ann Leow.

